# The Goldilocks Principle conferred by LDH isoenzymes controls murine T cell glycolysis and differentiation

**DOI:** 10.1101/2022.01.21.477227

**Authors:** Xuyong Chen, Lingling Liu, Ruoning Wang

## Abstract

Isoenzyme divergence is a prevalent mechanism governing tissue-specific and developmental stage-specific metabolism in mammals. The isoenzyme pattern (spectrum) of lactate dehydrogenase (LDH) reflects the status of glucose metabolism in different organs such as muscle, liver, and heart. T cells are highly dependent on glucose metabolism for survival, proliferation, and differentiation. However, the LDH isoenzyme spectrum in T cells and its potential impact on T cell-mediated immune response remain unclear. Here, we discovered that the LDH spectrum in murine T cells is characterized by three tetrameric isoenzymes composed of LDHA and LDHB (LDH-3/4/5). Genetic deletion of LDHA or LDHB altered the isoenzyme spectrum by removing all heterotetramers and leaving T cell with LDH-1 (the homotetramer of LDHB) or LDH-5 (the homotetramer of LDHA), respectively. Accordingly, altering the isoenzyme spectrum by deleting LDHA or LDHB compromises T cell metabolic fitness and effector cell differentiation in vivo. Unexpectedly, deleting LDHA suppressed glycolysis, whereas deleting LDHB further enhanced it, indicating that an optimal zone of glycolytic activity is critical to driving effector T cell differentiation. Mechanistically, the LDH isoenzyme spectrum imposed by LDHA and LDHB is required to optimize glycolysis to maintain a balanced NAD^+^/NADH pool, a hallmark of metabolic fitness. Together, our results suggest that the LDH isoenzyme spectrum enables “Goldilocks levels” of glycolytic and redox activity (i.e., not too high, not too low, but just right) to control T cell differentiation function.

## Introduction

Gene duplication drives the evolution of isoenzymes that differ in polypeptide sequences, quaternary structures, and catalytic efficiencies but catalyze the same biochemical reactions^1^. The species-specific features of isoenzymes may reflect the biochemical adaption of the species to environmental conditions^2^. Within a single organism, the divergence of isoenzyme pattern (spectrum) fine-tunes metabolic activities to meet the need of a specific tissue or developmental stage. T cell-dependent immune response is energetically costly and requires an optimal allocation of glucose-derived carbon to support rapid expansion and differentiation. It is conceivable that the isoenzyme spectrum plays a critical role in shaping T cell metabolic profile, which is characterized by high aerobic glycolysis, increased glutaminolysis but decreased fatty acid β-oxidation after activation ^3–5^. Lactate dehydrogenase (LDH) controls the last step of glycolysis by catalyzing the coupled interconversion of pyruvate to lactate and NADH to NAD^+^. Besides allosteric regulation, substrate-level regulation, and gene expression regulation, LDH is known to have predominantly five isoenzymes (LDH-1 to LDH-5), each of which has distinct catalytic properties ^6^. The LDH isoenzyme spectrum reflects glucose metabolism status in different organs such as muscle, liver, and heart. However, the LDH isoenzyme spectrum in T cells and its potential impact on T cell-mediated immune response remain unclear. Therefore, we sought to examine the T cell-specific LDH isoenzyme spectrum in mice.

## Results

### Distinct isoenzyme spectrum characterizes T cell at different stages

The zymography of LDH revealed that the thymus exhibits a similar LDH isoenzyme spectrum as the heart with nearly all five isoenzymes present; however, the spleen and the lymph node have one dominant isoenzyme (LDH-5) and two minor isoenzymes (LDH-4 and LDH-3) (**Fig. 1A**). To assess the isoenzyme spectrum in T cells, we isolated T cells from various lymphoid organs and exam their zymography pattern. Similar to the pattern revealed in tissue, T cells in the thymus exhibit five isoenzymes, whereas T cells in the spleen and the lymph node only have LDH-3, LDH-4, and LDH-5 (**Fig. 1B**). LDH isoenzymes are homotetramers or heterotetramers of the translational products of the LDHA gene and the LDHB gene ^6^. TCR activation increased LDHA but reduced LDHB expression, accompanied by a moderate reduction of LDH-3 and LDH-4 intensity (**Fig. 1C, E and G**). Next, we sought to assess the role of each gene product in regulating the LDH isoenzyme spectrum and enzymatic activities. Specifically, we generated T cell-specific *LDHA*, *LDHB*, and *LDHA/B* double knockout strains (*LDHA*, *LDHB*, *LDHA/B* dKO) by crossing the CD4-Cre strain with the *LDHA^fl^* strain and *LDHB^fl^* strain separately or consecutively. qPCR and immunoblot analysis validated the deletion of each gene and protein (**Fig. 1F and H**). Accordingly, deleting LDHA leads to the loss of LDH-3 through LDH-5, but the formation of LDHB homotetramer isoenzyme LDH-1. Conversely, deleting LDHB leads to the loss of LDH-3 and LDH-4 without affecting LDH-5, the LDHA homotetramer isoenzyme. Importantly, deleting both genes abolishes all the isoenzymes (**Fig. 1D**). Notably, the remaining weak stain in the LDH-5 position in the LDHA cKO or LDHA/B dKO samples is likely the sample loading background, which often overlaps with LDH-5 since LDH-5 has the slowest electrophoretic mobility among all the LDH isoenzymes. Interestingly, alteration of LDH isoenzyme spectrum due to LDH gene deletion did not result in any defects in T cell development in the thymus, spleen, and lymph nodes (**Fig. S1A-B**). We demonstrated that transcriptional factors Myc and HIF-1α control glycolysis during T cell activation and differentiation by regulating metabolic gene expression^7,8^. However, neither Myc deletion nor HIF-1α deletion changes the LDH isoenzyme spectrum in T cells (**Fig. S2A-B**), indicating that the intrinsic properties of LDHA and LDHB proteins determine the isoenzyme spectrum. Collectively, our results show that thymic and periphery T cells exhibit distinct isoenzyme spectra. However, alteration of the LDH isoenzyme spectrum does not affect T cell development.

**Figure 1.**
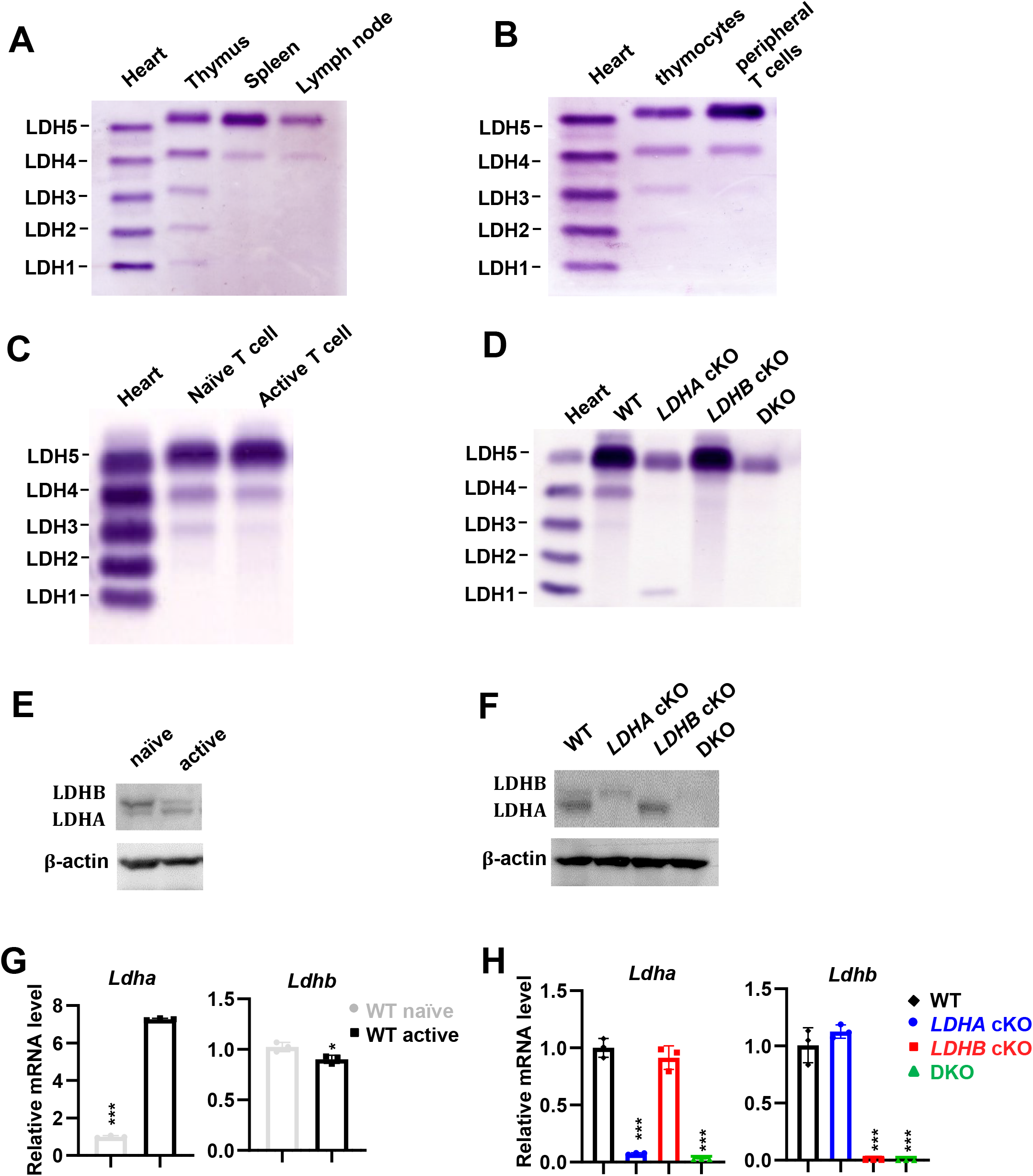
The LDH isoenzyme spectrum in T cells is characterized by distinct homotetramer and heterotetramers comprising LDHA and LDHB. (**A-D**) The LDH isoenzyme spectrum of the indicated tissues and cells was determined using the QuickGel system by zymography. (**E-F**) The protein level of LDHA and LDHB of the indicated cells was determined by immunoblot. (**G-H**) The relative mRNA levels of Ldha and Ldhb of the indicated groups were measured by qPCR. Data represent mean±SEM (n=3) for each group, experiments were repeated for 2 times, \**p* < 0.05, \*\**p* < 0.01, \*\*\**p* < 0.001, Student’s t-test (**G**), one-way ANOVA (**H**).

### The optimized LDH isoenzyme spectrum is required to sustain T cell expansion and inflammation in vivo

To assess the impact of altering the LDH isoenzyme spectrum on CD4^+^ T cells in vivo, we first employed a well-established competitive homeostatic proliferation assay to determine the ratio and carboxyfluorescein succinimidyl ester (CFSE) dilution pattern of purified WT(*Thy1.1*^+^), *LDHA cKO*, *LDHB cKO or LDHA/B dKO*(*Thy1.2*^+^) CD4^+^ T cells in a lymphopenic host (Rag^−/−^). Deleting both LDHA and LDHB genes loses all five isoenzymes, almost abolishing cell proliferation and viability as evidenced by the cell ratio between dKO and WT and CFSE dilution pattern (**Fig. 2A-C**). Importantly, altering the isoenzyme spectrum by deleting either LDHA alone or LDHB alone is sufficient to dampen cell proliferation and viability. However, LDHA cKO cells exhibit more profound defects than LDHB cKO cells (**Fig. 2A-C**). Next, we sought to measure antigen-specific, TCR-dependent proliferation and differentiation of T cells in the context of modulating the LDH isoenzyme spectrum. Since dKO completely abolishes the LDH isoenzyme spectrum, we focused on LDHA or LDHB cKO in the experiment. We crossed *Thy1.1* and CD4-Cre, *LDHA^fl^ or LDHB^fl^* mice with OT-II transgenic mice to generate OT-II, WT(*Thy1.1*^+^), OT-II, *LDHA cKO*(*Thy1.2*^+^) and OT-II, *LDHB cKO*(*Thy1.2*^+^) donor strains in CD45.2^+^ background. We then adoptively transferred mixed, and CFSE labeled WT and cKO CD4^+^ T cells into CD45.1^+^ mice immunized with chicken ovalbumin (OVA_323-339_). Consistent with the homeostatic proliferation results, altering the isoenzyme spectrum by deleting either LDHA or LDHB led to proliferation defects in an antigen-specific manner after immunization. Also, LDHA cKO cells have more severe proliferation defects than LDHB cKO cells (**Fig. 2D-F**). Interestingly, LDHA or LDHB deletion resulted in less IFNγ^+^CD4^+^ T cells and IL17^+^CD4^+^ T cells than the WT group, indicating that the alteration of the isoenzyme spectrum influences T cell differentiation (**Fig. 2G and S3A-B**). Experimental autoimmune encephalomyelitis (EAE) is an inflammatory demyelinating disease mouse model and is primarily driven by the proinflammatory T cells (T helper 1/T_H_1 and T helper 17/T_H_17) in the central nervous system. A recent study has shown that the genetic deletion of LDHA in T cells conferred protection against the pathogenic progression of EAE ^9^. Next, we used this well-characterized system to investigate the in vivo CD4^+^ T cell response in the context of altering the LDH isoenzyme spectrum by deleting the LDHB gene. In line with our homeostatic and antigen-specific proliferation results, deleting LDHB in T cells dramatically reduced CD4^+^ T cell infiltration in the CNS and slowed the pathogenic progression of mice (**Fig. 2H and S3C**). Collectively, these findings suggest that the optimal LDH isoenzyme spectrum conferred by LDHA and LDHB is required to sustain T cell expansion and differentiation in vivo. Thus, altering the isoenzyme spectrum by deleting either LDHA or LDHB dampens the metabolic fitness of T cells and hence causes proliferation and differentiation defects in vivo.

**Figure 2.**
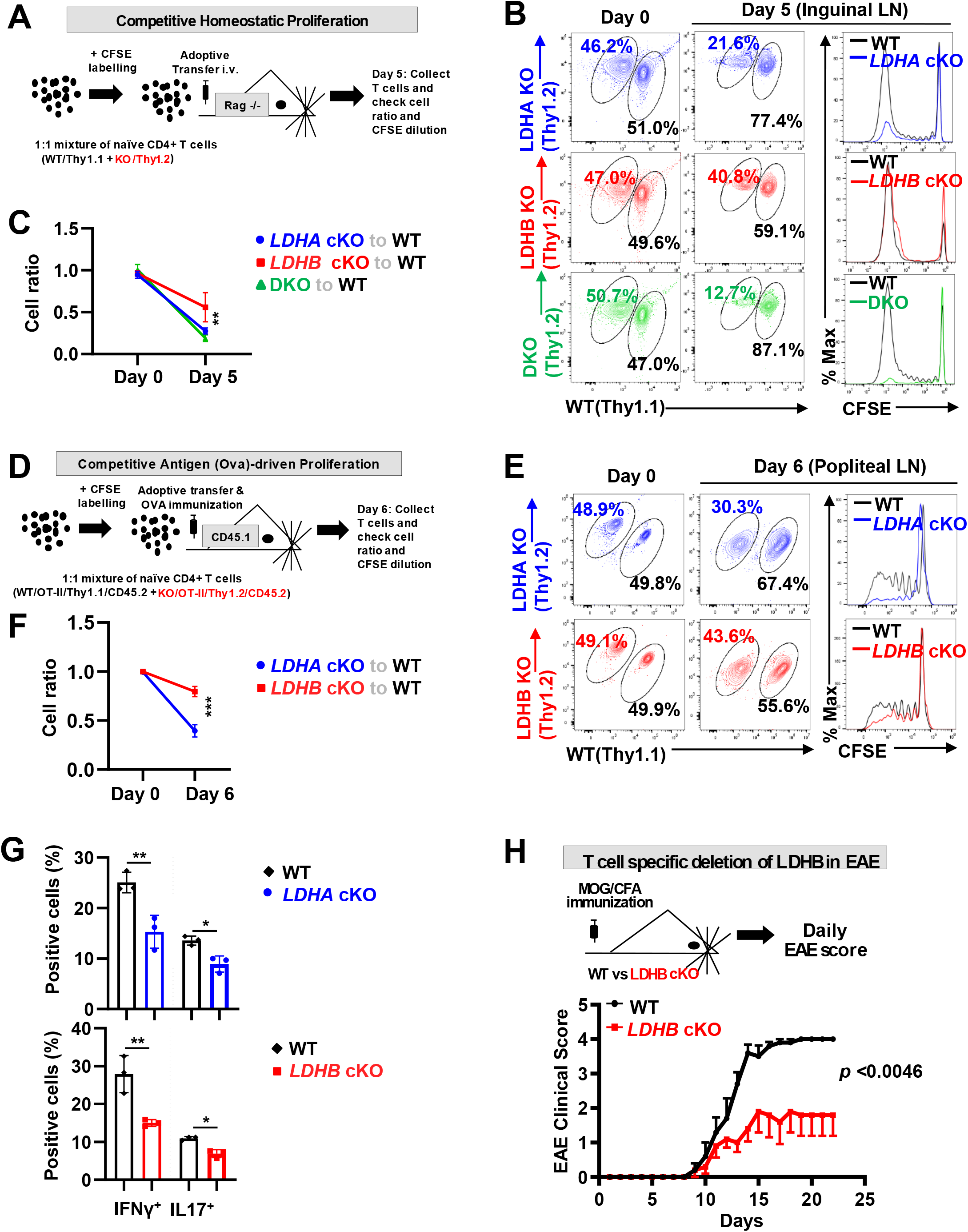
The optimized isoenzyme spectrum requires LDHA and LDHB to sustain T cell expansion and inflammation in vivo. (**A&D**) Diagrams of the indicated experimental procedures. **(B-G)** The donor CD4^+^ T cell ratios before and after adoptive transfer were determined by surface staining of isogenic markers. Cell proliferation was determined by CFSE dilution. The indicated cytokines were determined by intracellular staining. (**H**) Diagram of EAE experimental procedure (top panel), clinical scores were evaluated daily (bottom panel), data represent mean±SEM (n=3-5) for each group, experiments were repeated for 2 times, \*\**p* < 0.005, \*\*\**p* < 0.001, two-way ANOVA.

### Alteration of the LDH isoenzyme spectrum impacts T cell proliferation and differentiation in vitro

Next, we employed a range of in vitro functional assays to investigate how altering the LDH isoenzyme spectrum impacts T cell phenotypes. We reasoned that a proximately physiological relevant oxygen level (around 5% O_2_) supplied in vitro might resemble the effects of modulating LDH and glycolysis on T cells functions in vivo ^10^. Hence, we performed in vitro cell culture studies under 5% O_2_. In contrast to a dispensable role of LDHB in regulating T cell activation, LDHA and LDHA/B deletion caused more cell death (**Fig.S4 A-B**) and delayed cell cycle progression from G0/1 to S phase (**Fig.S4C**) after CD4^+^ T cells activation in vitro, which were associated with reduced cell surface activation markers (**Fig. S4D**), moderately reduced cell size (**Fig. S4E**) and RNA/DNA/Protein contents (**Fig. S4F-G**). Next, we mixed CFSE labeled WT(*Thy1.1*^+^) CD4^+^ T cells with *LDHA cKO*, *LDHB cKO*, or *LDHA/B dKO*(*Thy1.2*^+^) CD4^+^ T cells at 1:1 ratio and followed cell ratio and proliferation in the competitive setting after activation. In line with the in vivo results, altering the LDH isoenzyme spectrum by deleting LDHA, LDHB, or both genes significantly reduced the cell number and the ratio of KO to WT in a time-dependent manner (**Fig.3 A-C**). Deleting a single gene moderately affects CFSE dilution, while deleting both genes in CD4^+^ T cells dramatically reduced CFSE dilution compared to WT cells (**Fig.3 D**). Next, we examined the effects of altering the LDH isoenzyme spectrum on T_H_1 and T_H_17 differentiation. In line with the OT-II and EAE results, altering the isoenzyme spectrum due to LDHB deletion reduced the percentage of IFNγ^+^CD4^+^ T cells IL17^+^CD4^+^ T cells under T_H_1 and T_H_17 polarization conditions, respectively (**Fig.3 F**). Consistent with the recent studies^9 11^, LDHA deletion significantly suppressed T_H_1 and T_H_17 differentiation (**Fig.3 E**). Similarly, the complete abrogation of the LDH activity by deleting both LDHA and LDHB resulted in defects in T_H_1 and T_H_17 differentiation (**Fig.3 G**). Collectively, in vitro findings corroborate with in vivo results, suggesting that the optimal LDH isoenzyme spectrum is required to sustain T cell expansion and differentiation.

**Figure 3.**
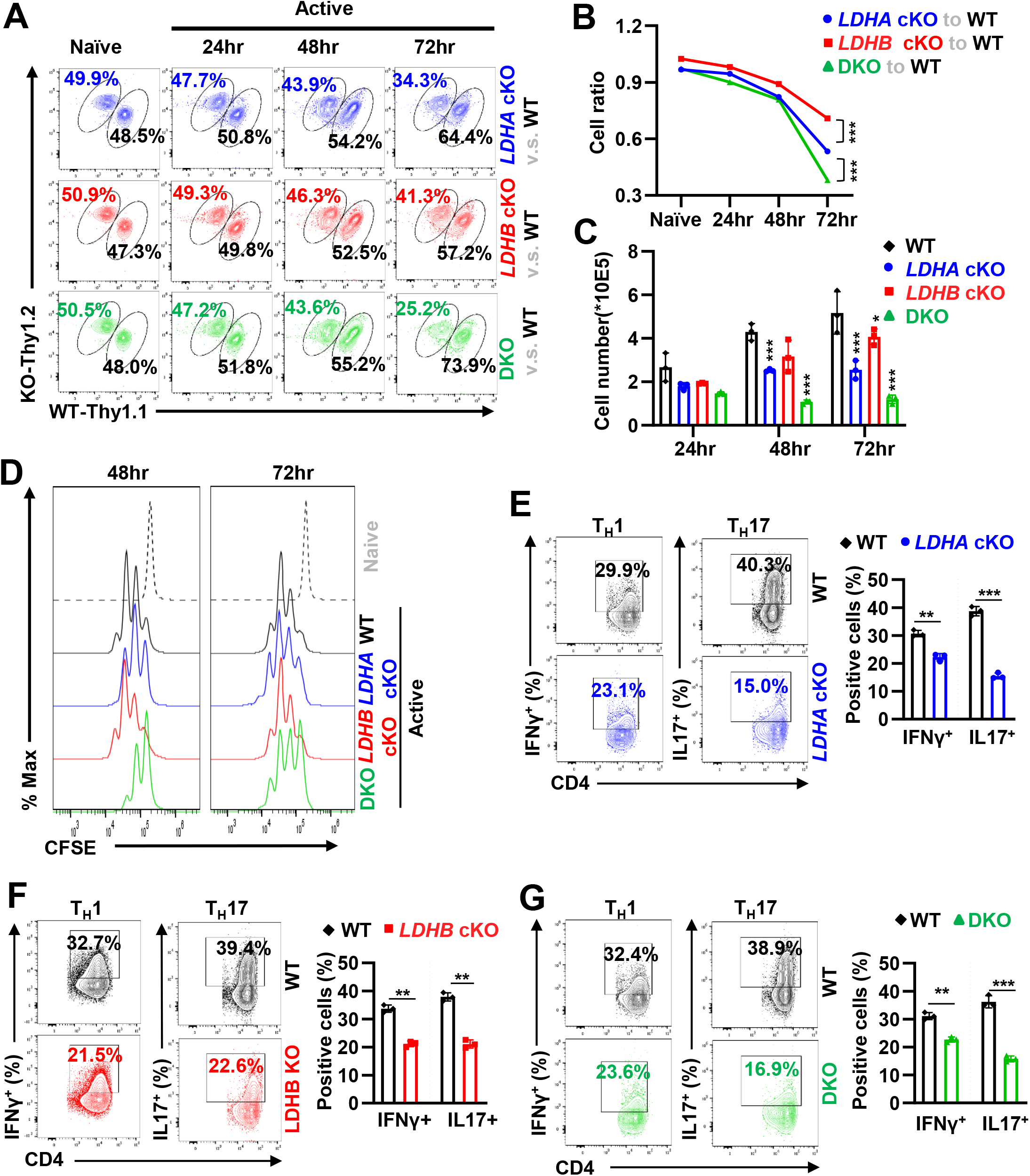
Alteration of the LDH isoenzyme spectrum impacts T cell proliferation and differentiation in vitro. **(A-D)** The CD4^+^ T cells with indicated genotypes were mixed with a 1:1 ratio and activated in vitro. At the indicated time points, cell ratio was determined by surface staining of isogenic markers **(A and B)**; cell number was determined by the cell counter **(C)**, and cell proliferation was determined by CFSE dilution **(D)**. Data represent of 3 independent experiments, \**p* < 0.05, \*\*\**p* < 0.001, one-way ANOVA. **(E-G)** CD4^+^T cells with indicated genotypes were polarized toward T_H_1 and T_H_17 lineages for 72 hours. The indicated proteins were quantified by intracellular staining. Data represent of 3 independent experiments, \*\**p*<0.01, \*\*\**p*<0.001, Student’s *t* test.

### The Goldilocks Principle conferred by the LDH isoenzyme spectrum enables a fine-tuned redox control of T cell differentiation

We then sought to gain more mechanistic insights on how altering the LDH isoenzyme spectrum impacts T cell metabolism and differentiation. We and others have previously shown that T_H_17 cell sustains high glycolysis, and inhibiting glycolysis suppresses T_H_17 cell differentiation ^8 12,13^. We reasoned that altering the LDH isoenzyme spectrum by LDHA or LDHB deletion would reduce glycolytic activities and suppress TH1 and T_H_17 differentiation. Indeed, deleting LDHA suppresses glucose consumption, lactate production, extracellular acidification rates (ECAR), and glycolysis (^3^H_2_O production from [5-^3^H]-glucose at the triose phosphate isomerase reaction) (**Fig. 4A-D**). Surprisingly, deleting LDHB resulted in an opposite pattern of changes in the above metabolic assays (**Fig. 4A-D**). Specifically, altering the LDH isoenzyme spectrum by LDHB deletion, compared to WT control, does not suppress but enhance the overall glycolysis flux as indicated by an increased glucose consumption and lactate production (**Fig. 4A-B**), a higher EACR (**Fig. 4C**), and a heightened ^3^H_2_O production from [5-^3^H]-glucose (**Fig. 4D**) ^14^. However, LDHB deletion did not change the overall glutamine consumption, glutamate production, glutaminolysis, and fatty acid β-oxidation (FAO) (**Fig. S5 A-D**). Also, LDHB deletion did not alter these pathways’ metabolic gene expression profile (**Fig. S5E-G**). The ratio of free cytosolic [NAD^+^]/[NADH] is fundamentally crucial in ensuring cellular redox and energy homeostasis^15^. Glycolysis determines the redox state of NAD^+^/NADH in the cytoplasm through the action of GAPDH and LDH ^16^. Next, we measured the cellular NAD^+^/NADH ratio after altering the LDH isoenzyme spectrum. LDHA deletion or LDHB deletion in T cells decreased or increased the NAD^+^/NADH ratio, respectively (**Fig. 4E**). To test if the imbalance of NAD^+^/NADH causes differentiation defects in LDHA and LDHB cKO cells, we added NADH to LDHB cKO cells and NAD^+^ to LDHA cKO cells under T_H_17 and T_H_1 polarization conditions. Remarkably, NADH supplement and NAD^+^ supplement restored the balance of NAD^+^/NADH, and partially rescued differentiation defects in LDHB cKO and LDHA cKO, respectively (**Fig. 4F, G, I, J and S6A, B, D, E**). Notably, altering the LDH isoenzyme spectrum also caused the reduction of cellular ATP level, which was restored after NAD^+^ or NADH supplement in LDHA cKO and LDHB cKO cells (**Fig. 4H, K and S6C, F**). These findings suggest that the LDH isoenzyme spectrum is required to maintain the redox state of NAD^+^/NADH in T cells. Collectively, our results further indicate the presence of the ‘‘Goldilocks zone” (not too high, not too low, but just right) may ensure the optimal glucose carbon input in degerming the immunological outcomes quantitively (**Fig. 5**).

**Figure 4.**
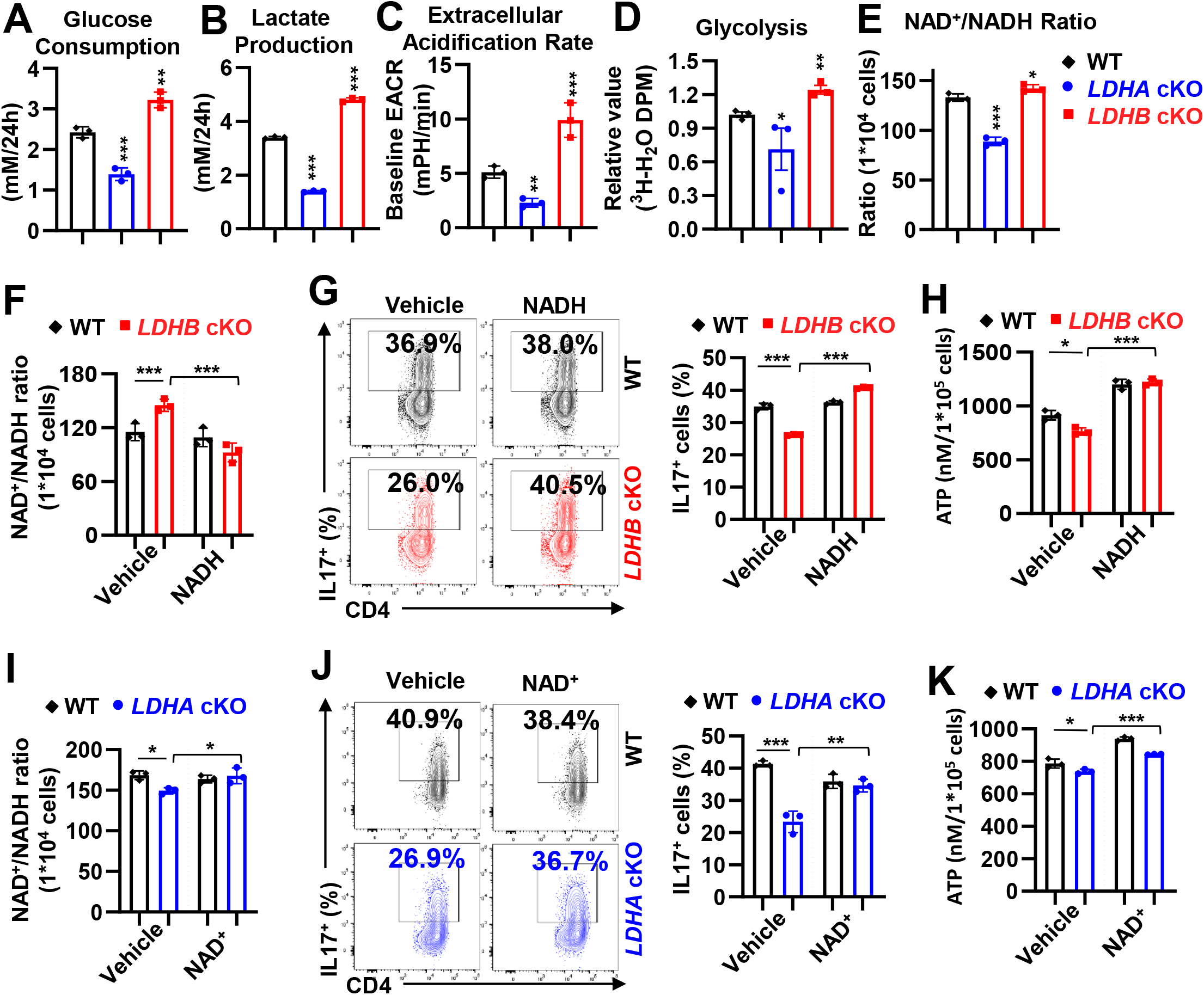
LDH isoenzyme spectrum confers an optimal glycolytic flux to maintain the NAD^+^/NADH ratio during T cell differentiation. **(A-B)** CD4^+^T cells with indicated genotypes were polarized toward T_H_17 lineages for 2 days and then were resuspended with fresh media. After 24 h, blank (without cells) media and spent (with cells) media were collected. The indicated metabolites were quantified by the bioanalyzer (YSI 2900). The consumption and production of indicated metabolites were determined by the difference between blank and spent media. (**C-K**) CD4^+^T cells with indicated genotypes were polarized toward T_H_17 lineages for 3 days, the extracellular acidification rate (ECAR) was measured by Seahorse XF96 Analyzer (**C**). Glycolysis rate was determined by generating ^3^H_2_O) from [5-^3^H]-glucose (**D**). NAD^+^/NADH ratio was measured by the NAD/NADH-Glo^™^ Assay kit (**E-F, and I**). T_H_17 differentiation was determined by the intracellular staining of IL-17 (**G and J**), ATP levels were determined by the CellTiter-Glo^®^ 2.0 Assay kit (**H and K**). Data represent of 3 independent experiments, \**p* < 0.05, \*\**p* < 0.01, \*\*\**p* < 0.001, one-way ANOVA.

**Figure 5.**
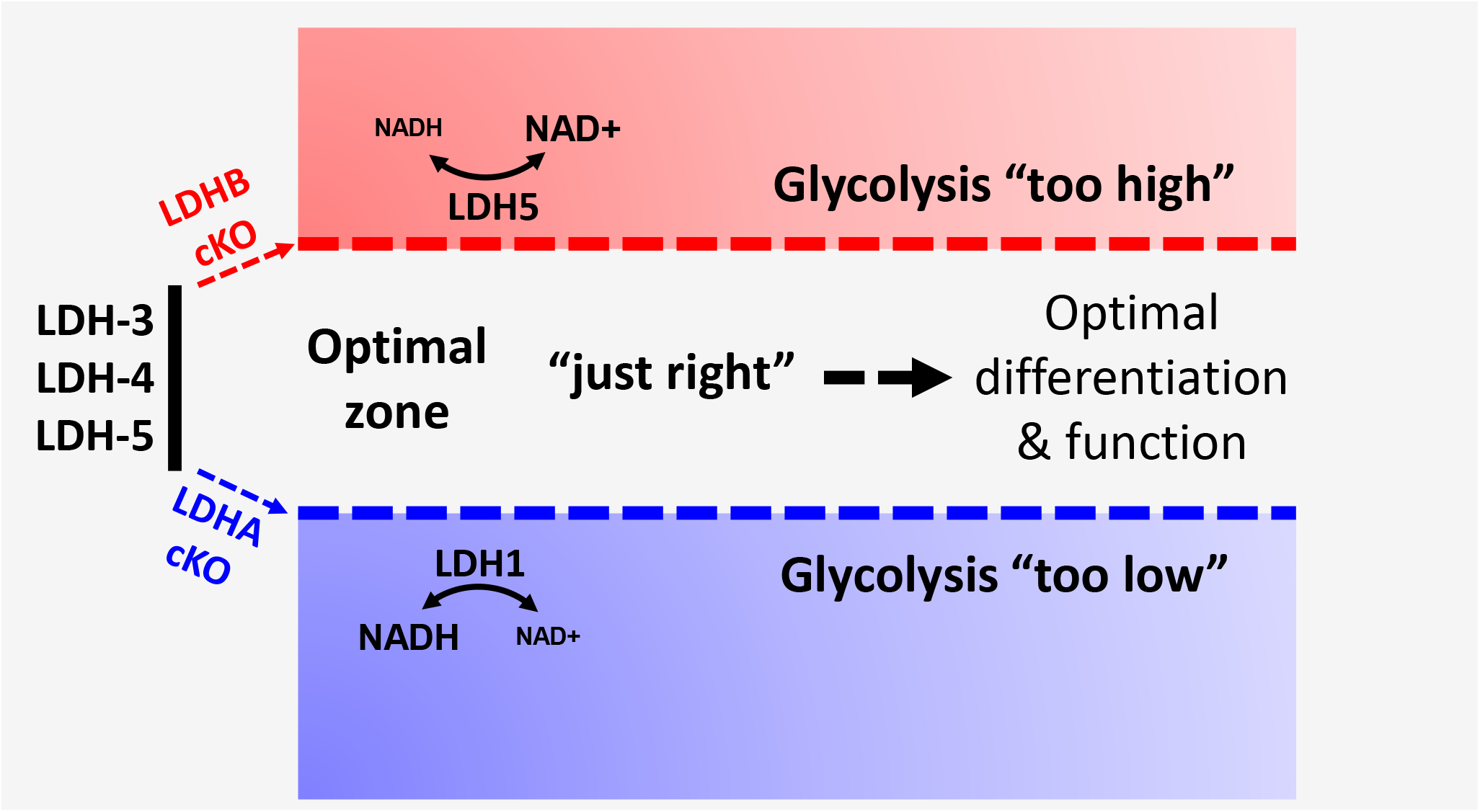
The ‘‘Goldilocks principle” in regulating glycolysis and differentiation. Proposed conceptual model: the LDH isoenzyme spectrum enables “Goldilocks levels” of glycolytic and redox activity (i.e., not too high, not too low, but just right) to control T cell differentiation function.

## Discussion

T cell-mediated immune response in the vertebrate is energetically costly and often requires rapid employment and allocation of available carbon resources. Hence, changing environments and rapidly evolving pathogens impose selective pressures on T cells, ensuring rapid metabolic changes upon activation. Accordingly, TCR stimulation leads to an immediate engagement of aerobic glycolysis, transforming carbon and chemical energy to support T cell growth, proliferation, and differentiation^14,17–21^. Remarkably, one of the largest constituents of the T cell proteome is the reservoir of highly abundant glycolytic enzymes, including LDHA and LDHB ^22–24^. Without changing transcriptional and translational states significantly, a diverse and broad spectrum of LDH isoenzymes in T cells may permit a rapid glycolytic switch upon activation while maintaining some degrees of flexibility to optimize activities across a wide range of nutrient conditions and immunological contexts^7,8,12,25^. The five LDH isoenzymes (LDH-1 to 5) are tetrameric proteins arising from varying combinations of LDHA and LDHB. The LDH isoenzyme spectrum in central organs such as muscle, liver, and heart reflect their metabolic profiles, particularly the status of glycolysis. We have revealed that the LDH isoenzyme spectrum in periphery T cells consists of one homotetramer of LDHA (LDH-5) and two heterotetramers of LDHA and LDHB (LDH-3 and LDH-4). Despite their apparent redundancy, removing heterotetramers (LDH-3 and LDH-4) and leaving T cells with either LDH-1 (LDHA cKO) or LDH-5 (LDHB cKO) compromised T cell survival and differentiation significantly, suggesting that a diverse spectrum of the LDH isoenzyme confers the optimal metabolic fitness to T cell.

In the final step of glycolysis, LDH catalyzes the reversible conversion of pyruvate to lactate, coupled with the interconversion of NADH to NAD^+^. Abrogating LDHA shifts the isoenzyme spectrum towards the LDH-1, reducing the overall glycolytic flux and decreasing effector T cell proliferation and differentiation. Remarkably, shifting the LDH isoenzyme spectrum toward LDH-5 by LDHB deletion enhances glycolysis but still compromises T cell survival and differentiation. One important principle of biological processes is to avoid reaching the extreme but evolve toward optimization ^26^. Our findings suggest that glycolytic flux in T cells needs to be managed within certain margins, “not too high, not too low, but just right” (often called the Goldilocks principle), to ensure the optimal functional outcome in T cells. Similarly, the optimal strength of TCR signaling governs how T cells transition between different states during development in the thymus, activation, and differentiation in the periphery ^27,28^. The Goldilocks zone of glycolysis reflects T cell’s capacity to ensure proper functions with high metabolic efficiency. However, it is conceivable that the boundaries of the Goldilocks zone may change as T cells transition between different states during activation and differentiation. Supporting this idea, different T cell subsets exhibit distinct glycolytic capacities ^8,12,29^.

Our studies revealed that altering one glycolytic enzyme’s isoenzyme spectrum is sufficient to tune up or down the overall glycolysis flux in T cells. The LDH activity is not only indispensable for sustaining glycolytic flux but also is critical for controlling the ratio of free cytosolic [NAD^+^]/[NADH], which reflects the overall redox state in hyperproliferating cells such as active T cells and cancer cells ^15,16,30–33^. Since each isoenzyme exhibits a different cofactor/substrate affinity, the composition of multiple LDH isoenzymes in T cells reflects an intrinsic constraint on keeping the glycolytic rate within a given range and keeping the NAD^+^/NADH ratio in check. Consequentially, the suboptimal glycolytic rate resulting from altering the isoenzyme spectrum leads to the imbalance of the NAD^+^/NADH pool. The redox state of NAD^+^/NADH determines cellular redox balance and thus controls cellular homeostasis and fate ^34,35^.

Glycolysis also impacts T cell redox homeostasis through other mechanisms. Glycolysis interconnects with the pentose phosphate pathway and, therefore, may influence the cytosolic ratio of NADP^+^/NADPH, the other cellular redox couple. Through channeling its intermediate metabolite to glycine biosynthesis, glycolysis also contributes to the production of glutathione, an essential antioxidant. The most critical role of LDH-dependent reaction is to preserve cytosolic NAD^+^/NADH homeostasis under various cellular contexts ^36,37^. Supporting this idea, we have shown that restoring the ratio of NAD^+^/NADH in LDH-1 dominant (LDHA cKO) T cells by NAD^+^ supplement or in LDH-5 dominant (LDHB cKO) T cells by NADH supplement could partially rescue differentiation defects in these cells. Collectively, the LDH isoenzyme spectrum may evolve to ensure the optimal range of glycolytic rate to maintain redox balance and support T cell function across a broad range of nutrient conditions and immunological contexts.

## Data availability statement

The RNA-seq datasets generated for this study can be found in the GEO accession GSE190799.

## Author contributions

R. Wang conceptualized and supervised this work. X.C. carried out the experiments. X.C. and L.L. were involved in data collection, analysis, and review. X.C., and R. Wang wrote the manuscript. All authors discussed the results and provided feedback on the manuscript.

## Materials and Methods

### Mice

C57BL/6NHsd (WT) mice, Rag^−/−^ mice, OTII mice, Thy 1.1^+^ mice, CD45.1^+^ and Ldha^fl^ mice were obtained from Jackson Laboratory (Bar Harbor, ME). Ldhb^<tm1a((KOMP)Wtsi>^ mice was generated by UC Davis, KOMP repository and was further crossed with a transgenic Flippase strain (B6.129S4*Gt(ROSA)26Sor^tm1(FLP1)Dym^*/RarnJ) to remove the LacZ-reporter allele to generate LDHB^fl^ mice. LDHA^fl^ mice and LDHB^fl^ mice were crossed with the CD4-Cre strain to generate T cell-specific *LDHB* knockout strain (*LDHB* cKO) or *LDHA* knockout strain (*LDHA* cKO). The LDHA-LDHB double knockout (DKO) mice were generated by crossing LDHA cKO mice with LDHB cKO mice. *HIF1-α* cKO and *c-MYC* KO mice were reported previously ^7,8^. OT-II mice (B6.Cg-Tg(TcraTcrb)425Cbn/J) were crossed with CD4Cre *LDHA* cKO mice or CD4Cre *LDHB* cKO mice to generate the OT-II CD4Cre *LDHA* cKO mice or OT-II CD4Cre *LDHB* cKO mice. OT-II mice (B6.Cg-Tg(TcraTcrb)425Cbn/J) were crossed with *Thy1.1^+^* mice (B6.PL-*Thy1*a/CyJ) to generate OT-II *Thy1.1*(WT) mice. Mice with gender and age-matched (7-12 weeks old) were used in the experiments. All mice were bred and kept in specific pathogen-free conditions at the Animal Center of Abigail Wexner Research Institute at Nationwide Children Hospital. Animal protocols were approved by the Institutional Animal Care and Use Committee of the Abigail Wexner Research Institute at Nationwide Children’s Hospital (IACUC; protocol number AR13-00055).

### Mouse T cell isolation and culture

All cells were cultured in RPMI 1640 media supplemented with 10% (v/v) fetal bovine serum (FBS), 2 mM L-glutamine, 0.05 mM 2-mercaptoethanol, 100 units/mL penicillin and 100 μg/mL streptomycin at 37°C in 5% CO_2_ and 5% O_2_. Total CD3^+^ T cells or naïve CD4^+^ T cells were enriched from mouse spleen and lymph nodes by negative selection using MojoSort Mouse CD3 T Cell Isolation Kit or MojoSort Mouse CD4 Naive T Cell Isolation Kit (MojoSort, Biolegend) following the manufacturer’s instructions. For the activation assay, freshly isolated total T cells were either maintained in culture media with 5 ng/mL IL-7 for resting state or were activated with 5 ng/mL IL-2 and plate-bound anti-CD3 (clone 145-2C11) and anti-CD28 (clone 37.51). The culture plates were pre-coated with 2 μg/mL anti-mCD3 (clone 145-2C11, Bio X Cell) and 2 μg/mL anti-mCD28 (clone 37.51, Bio X Cell) antibodies overnight at 4°C. For CD4^+^ T cell differentiation, 48 wells culture plates were pre-coated with 2 μg/mL (T_H_1 differentiation) or 4 μg/mL (T_H_17 differentiation) anti-mCD3 and anti-mCD28 antibodies overnight at 4°C. Freshly isolated naïve CD4^+^ T cells (0.6 ×10^6^/mL) were activated with plate-bound antibodies and mIL-2 (200 U/mL), mIL-12 (5 ng/mL, PeproTech) (T_H_1 differentiation), or with mIL-6 (50 ng/mL, PeproTech), hTGF-β1 (2 ng/mL), anti–mIL-2 (8 μg/mL, Bio X Cell), anti–mIL-4 (8 μg/mL, Bio X Cell), and anti–mIFN-γ (8 μg/mL, Bio X Cell) (T_H_17 differentiation). In some experiments, 3-10 μM NADH or 0.1 μM NAD^+^ were added to cell culture media to restore intracellular NAD^+^/NADH ratio. Additional information of cytokines, antibodies, and chemicals were list in table 1.

### Flow Cytometry

For analyzing surface markers, cells were stained in PBS containing 2% (w/v) BSA and the appropriate antibodies from Biolegend. For analyzing intracellular cytokine IFN-γ and IL-17A, T cells were stimulated for 4 hours with eBioscience^™^ Cell Stimulation Cocktail (eBioscience) before being stained with cell-surface antibodies. Cells were then fixed and permeabilized using Foxp3 Fixation/Permeabilization solution according to the manufacturer’s instructions (eBioscience^™^). Cell proliferation was assessed by CFSE staining per the manufacturer’s instructions (Invitrogen). Cell viability was assessed by 7-AAD staining per the manufacturer’s instructions (Biolegend). For analyzing DNA/RNA content, cells were collected and stained with surface markers before being fixed with 4% paraformaldehyde for 30 min at 4°C, followed by a step of permeabilization with FoxP3 permeabilization solution (eBioscience). Cells were stained with 7AAD for 5 minutes and then stained with pyronin-Y (4ug/ml, PE) for 30 min before being analyzed by flow cytometer with PerCP channel for 7AAD (DNA) and PE channel for pyronin-Y (RNA). Protein synthesis assay kit (Item No.601100, Cayman) was used for analyzing protein content. Briefly, cells were incubated with O-propargyl-puromycin (OPP) for 1 hour, then were fixed and stained with 5 FAM-Azide staining solutions before being analyzed by flow cytometer with FITC channel. For analyzing cell cycle profile, cells were incubated with 10 μg/mL BrdU for 1 hour, followed by cell surface staining, fixation, and permeabilization according to Phase-Flow Alexa Fluro 647 BrdU Kit (Biolegend). Flow cytometry data were acquired on Novocyte (ACEA Biosciences) and were analyzed with FlowJo software (TreeStar). Additional information on antibodies and dyes was listed in table 2.

### LDH isoenzymes assay

The LDH isoenzymes assay was performed according to the manufacturer’s instructions of the QuickGel LD isoenzymes kit (HELENA Laboratories, Cat.No. 3338). Briefly, tissue or cell samples were homogenized and diluted in the cold Tris buffer (10mM, pH=7.4) for protein quantification. Samples (each with 1μg of protein) were loaded on the agarose gel and electrophoresed for 5min in the QuickGel chamber at 600 Volts. Then, the gel was removed from the chamber and stained with QuickGel LD isoenzyme reagent at 45°C for 20mins. The gel was washed with the de-staining solution for 10mins and then with water for 5mins before being dried in the chamber and scanned for analysis.

### NAD^+^/NADH detection assay

The NAD^+^/NADH ratios were performed according to the manufacturer’s instructions of the NAD/NADH-Glo^™^ Assay kit (Promega, Cat. No. G9071). Briefly, 1×10^4^ cells were suspended with 50uL PBS in 96-well white-walled tissue culture plates, an equal volume of NAD/NADH-Glo Detection Reagent was added to each well. Samples were incubated at room temperature for 30-60 min. The luminescence was measured by the Molecular Devices SpectraMax M2 Multilabel Microplate Reader (Marshall scientific), data were analyzed by the GraphPad Prism 9.0.

### ATP levels measurement

The ATP levels were measured according to the manufacturer’s instructions of the CellTiter-Glo^®^ 2.0 Assay kit (Promega, Cat. No. G9241). Briefly, 1×10^5^ cells were suspended with 50uL PBS in 96-well white-walled tissue culture plates, an equal volume of CellTiter-Glo 2.0 reagent was added to each well. The plate was put on a shaker for gentle mixing and incubated at room temperature for 30 minutes. The luminescence was measured by the Molecular Devices SpectraMax M2 Multilabel Microplate Reader (Marshall scientific), data were analyzed by the GraphPad Prism 9.0.

### Glucose, lactate, glutamine, and glutamate quantification

The naïve CD4^+^ T cells were polarized under T_H_17 condition for 2 days before being collected and re-suspended in fresh media at the density of 2×10^6^/mL. Blank media (without cells) and spent media were collected after 24 hours. The levels of metabolites were measured by the bioanalyzer (YSI 2900). The metabolite consumption and production were determined by calculating the difference between blank and spent media.

### EACR measurement

ECAR was determined by seahorse XFe96 Analyzer (Agilent Technologies) according to the manufacturer’s manual. Briefly, 1.5×10^5^ cells were suspended in Seahorse XF RPMI Assay Media (with 2mM glucose and 2mM glutamine) and seeded in a poly D Lysine precoated XF96 microplates. The plate was centrifuged to immobilize cells and kept in a non-CO2 incubator for 30 min. Basal ECAR was measured on a seahorse XFe96 Analyzer (Agilent Technologies).

### Western blot analysis

For protein extraction, cells were harvested, lysed, and sonicated at 4 °C in a lysis buffer (50 mM Tris-HCl, pH 7.4, 150 mM NaCl, 0.5% SDS, 5 mM sodium pyrophosphate, protease, and phosphatase inhibitor tablet). Cell lysates were centrifuged at 13,000×g for 15 min, and the supernatant was recovered. The protein concentrations were determined using the Pierce^™^ BCA Protein Assay kit (Thermo Fisher Scientific).

The samples were boiled in NuPAGE^®^ LDS Sample Buffer and Reducing solution (Thermo Fisher Scientific) for 5 min. The proteins were separated by NuPAGE 4-12% Protein Gels (Thermo Fisher Scientific), transferred to PVDF membranes by using the iBlot Gel Transfer Device (Thermo Fisher Scientific), then incubated with primary antibodies (Table 3), followed by incubating with the secondary antibodies conjugated with horseradish peroxidase. Immunoblots were developed on films using the enhanced chemiluminescence technique.

### RNA extraction, RT-qPCR, and RNAseq

RNeasy Mini Kit (Qiagen) was used for RNA isolation. Random hexamers and M-MLV Reverse Transcriptase (Invitrogen) was used for cDNA synthesis. BIO-RAD CFX96^™^ Real-Time PCR Detection System was used for SYBR green-based quantitative PCR. The relative gene expression was determined by the comparative *CT* method, also referred to as the 2^−ΔΔ*CT*^ method. The data were presented as the fold change in gene expression normalized to an internal reference gene (beta2-microglobulin) and relative to the control (the first sample in the group). Fold change = 2 ^−ΔΔ*C*^_T_ = [(*CT* _gene of interest_-*CT* _internal reference_)] sample A - [(*CT* _gene of interest_-*CT* _internal reference_)] sample B. Samples for each experimental condition were run in triplicated PCR reactions. Primers were used for detecting target genes (Table 4).

For RNA sequencing analysis, total RNA was extracted using RNeasy Mini Kit (Qiagen) and treated with DNase I according to the manufacturer’s instructions. After assessing the quality of total RNA using an Agilent 2100 Bioanalyzer and RNA Nanochip (Agilent Technologies), 150 ng total RNA was treated to deplete the levels of ribosomal RNA (rRNA) using target-specific oligos combined with rRNA removal beads. Following rRNA removal, mRNA was fragmented and converted into double-stranded cDNA. Adaptor-ligated cDNA was amplified by limit cycle PCR. After library quality was determined via Agilent 4200 TapeStation and quantified by KAPA qPCR, approximately 60 million paired-end 150 bp sequence reads were generated on the Illumina HiSeq 4000 platform. Quality control and adapter trimming were accomplished using the FastQC (version 0.11.3) and Trim Galore (version 0.4.0) software packages. Trimmed reads were mapped to the Genome Reference Consortium GRCm38 (mm10) murine genome assembly using TopHat2 (version 2.1.0), and feature counts were generated using HTSeq (version 0.6.1). Statistical analysis for differential expression was performed using the DESeq2 package (version 1.16.1) in R, with the default Benjamini-Hochberg *p-value* adjustment method. The heatmap was generated using Graphpad prism 9.0 software to show the differential expression of the selected genes.

### Radioactive tracer-based metabolic assays

The metabolic assays were performed as described previously ^38^. Glycolysis was measured by the generation of ^3^H_2_O from [5-^3^H(N)] D-glucose, fatty acid oxidation was measured by the generation of ^3^H_2_O from [9,10-^3^H] palmitic acid, glutamine oxidation activity was measured by the generation of ^14^CO_2_ from [U-^14^C]-glutamine. For catabolic activities generation of ^14^CO_2_, five million T cells in 0.5 mL fresh media were dispensed into 7mL glass vials (TS-13028, Thermo) with a PCR tube containing 50μL 0.2N NaOH glued on the sidewall. After adding 0.5 μCi radioactive tracer, the vials were capped using a screw cap with a rubber septum (TS-12713, Thermo) and incubated at 37 *°C* for 2 hours. The assay was then stopped by injection of 100μL 5N HCL and kept vials at room temperate overnight to trap the ^14^CO_2_. The NaOH in the PCR tube was then transferred to scintillation vials containing 10mL scintillation solution for counting. A cell-free sample containing 0.5μCi [U-^14^C]-glutamine was included as a background control. For catabolic activities generating ^3^H_2_O), 1μCi radioactive tracer was added to the suspension of one million cells in 0.5mL fresh media in 48 wells, then incubated at 37 °C for 2 hours. The cells were transferred to a 1.5 mL microcentrifuge tube containing 50μL 5N HCL, placed in 20 mL scintillation vials containing 0.5 mL water with the vials capped and sealed. ^3^H_2_O was separated from other radio-labeled metabolites by evaporation diffusion overnight at room temperature. A cell-free sample containing 1μCi radioactive tracer was included as a background control.

### Adoptive cell transfer assays

For homeostatic proliferation in lymphopenic *Rag*^-/-^ mice, naïve CD4^+^ T cells isolated from donor mice were mixed with WT and KO cells at a 1:1 ratio and labeled with CFSE. Approximately 1×10^7^ cells in 150 μL PBS were transferred via retro-orbital venous injection into 6-8 week-old gender-matched host mice. Mice were sacrificed after 5 days, and lymph nodes were extracted from host mice, then processed for surface staining and flow analysis.

For antigen-driven proliferation using OTII mice, naïve CD4^+^ T cells isolated from OTII/CD45.2 TCR transgenic donor mice were mixed with WT and KO cells at a 1:1 ratio and labeled with CFSE. Approximately 1×10^7^ cells in 150 μL PBS were transferred via retro-orbital venous injection into 6-8 week-old gender-matched CD45.1 host mice. Host mice were immunized subcutaneously in the hock area (10 μL each site) in both legs with 1 mg/mL OVA^323–339^ peptide (InvivoGen) emulsified with CFA (InvivoGen). After 6 days of antigen-driven proliferation, lymph organs were extracted from host mice then processed for surface staining, intracellular staining, and flow analysis.

### Experimental Autoimmune Encephalomyelitis (EAE)

For induced EAE, mice were immunized subcutaneously with 100 μg of myelin oligodendrocyte glycoprotein (MOG)_35–55_ peptide emulsified in complete Freund adjuvant (CFA), which was made from IFA(Difco) plus mycobacterium tuberculosis (Difco). Mice were i.p. injected with 200 ng of pertussis toxin (List Biological) on the day of immunization and 2 days later. The mice were observed daily for clinical signs and scored as described previously ^8^.

In some experiments, mice were euthanized when control mice reached the onset of symptoms. The CNS (brain and spinal cord), spleen, and peripheral lymph nodes were collected and mashed to make a single cell solution. The cell suspension was centrifuged on a 30%/70% Percoll gradient at 500*g* for 30 min to isolate mononuclear cells from the CNS, followed by cell surface staining and flow cytometric analysis as described above.

### Statistical analysis

Statistical analysis was conducted using the GraphPad Prism software version 9.0.0 (GraphPad Software, Inc.). *P* values were calculated with two-way ANOVA for the EAE experiments. Unpaired two-tail Student’s t-test, multiple comparisons of one/ two-way ANOVA, were used to assess differences in other experiments. *P* values smaller than 0.05 were considered significant, with *p* < 0.05, *p* < 0.01, and *p* < 0.001 indicated as *, **, and ***respectively.

