## Supplementary material for "The Goldilocks Principle conferred by LDH isoenzymes controls murine T cell glycolysis and differentiation": SI

**Supplemental Figure 1. LDH is dispensable for normal T cell development after the double-positive stage.**

**(A-B)** The distribution of indicated T cell subsets in the thymus, spleen, and peripheral lymph node (LN) were determined by cell surface markers staining (mean±SEM, n=3). Cell numbers were calculated by the cell counter.

**Supplemental Figure 2. Transcription factors HIF1-α and c-Myc are dispensable for regulating the LDH isoenzyme spectrum in T cells.**

(**A**) The LDH isoenzyme spectrum from the indicated tissues and cells was determined by zymography using the QuickGel system. (**B**) The protein level of HIF1-α and c-Myc of the indicated cells was determined by immunoblot.

**Supplemental Figure 3. Changes in the LDH isoenzyme spectrum impact T cell differentiation in vivo.**

**(A-B)** The indicated intracellular proteins from *in vivo* antigen-specific T cells were quantified by flow cytometry. **(C)** Cells were isolated from indicated organs. The indicated cell surface markers and cell numbers were quantified by flow cytometry, data represent mean±SEM (n=5) for each group, $*p<0.05$, one-way ANOVA. n.s., not significant.

**Supplemental Figure 4. Alteration of the LDH isoenzyme spectrum differentially impacts T cell activation in vitro.**

**(A-B)** At the indicated time points after activation in vitro, the cell viability was assessed by 7AAD staining. **(C)** The cell cycle profile of the indicated cells (48 h after activation) was determined by Brdu labeling and 7AAD staining. **(D-G)** Cell surface markers, size, RNA/DNA/protein contents of the indicated cells (24 h after activation) were determined by flow cytometry. Data represent of 3 independent experiments, $*p < 0.01$, $**p < 0.001$, one-way ANOVA.

**Supplemental Figure 5. Altering the isoenzyme spectrum by LDHB deletion does not affect glutamine and fatty acid catabolism.**

**(A-B)** $CD4^+$T cells with indicated genotypes were polarized toward $T_H17$ lineages for 2 days and then were resuspended with fresh media. After 24 h, blank (without cells) media and spent (with cells) media were collected. The indicated metabolites were quantified by the bioanalyzer (YSI 2900). The consumption and production of indicated metabolites were determined by the difference between blank and spent media. (**C-G**) $CD4^{+}$T cells with indicated genotypes were polarized toward $T_H17$ lineages for 3 days, FAO and glutaminolysis rate were determined by the generation of $^{3}H_2O$ from [9,10-$^{3}$H] Palmitic acid and the generation of $^{14}CO_2$ from [U-$^{14}$C]-glutamine, respectively **(C-D).** Data represent mean±SEM (n=3) for each group, **$p < 0.01$, ***$p < 0.001$, one-way ANOVA. The expression of indicated metabolic genes was determined by RNAseq, the heat map represents the log2 value of the mRNA counts (see color scale) (**E-G**).

**Supplemental Figure 6. LDH isoenzyme spectrum confers an optimal glycolytic flux to maintain the $NAD^{+}$/NADH ratio during T cell differentiation.**

(**A-F**) $CD4^{+}$T cells with indicated genotypes were polarized toward $T_H1$ lineages for 3 days. $NAD^{+}$/NADH ratio was measured by the NAD/NADH-Glo™ Assay kit (**A and D**). $T_H1$ differentiation was determined by the intracellular staining of IFN-γ (**B and E**), ATP levels were determined by the CellTiter-Glo® 2.0 Assay kit (**C and F**). Data represent of 3 independent experiments, *$p < 0.05$, **$p < 0.01$, ***$p < 0.001$, one-way ANOVA.
