## Supplementary material for "The Goldilocks Principle conferred by LDH isoenzymes controls murine T cell glycolysis and differentiation": S figures

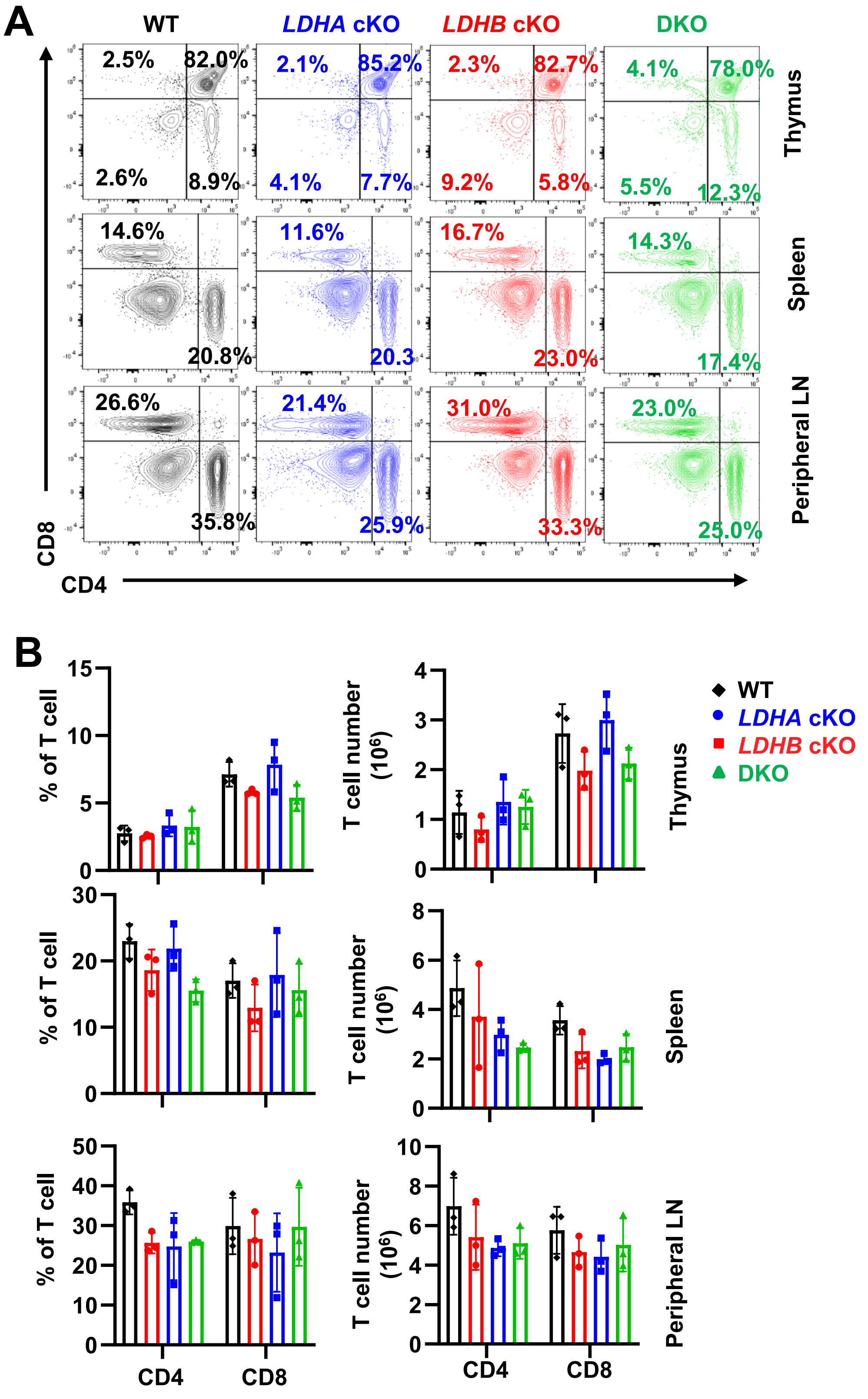


**Supplemental Figure 1. LDH is dispensable for normal T cell development after the double-positive stage.**

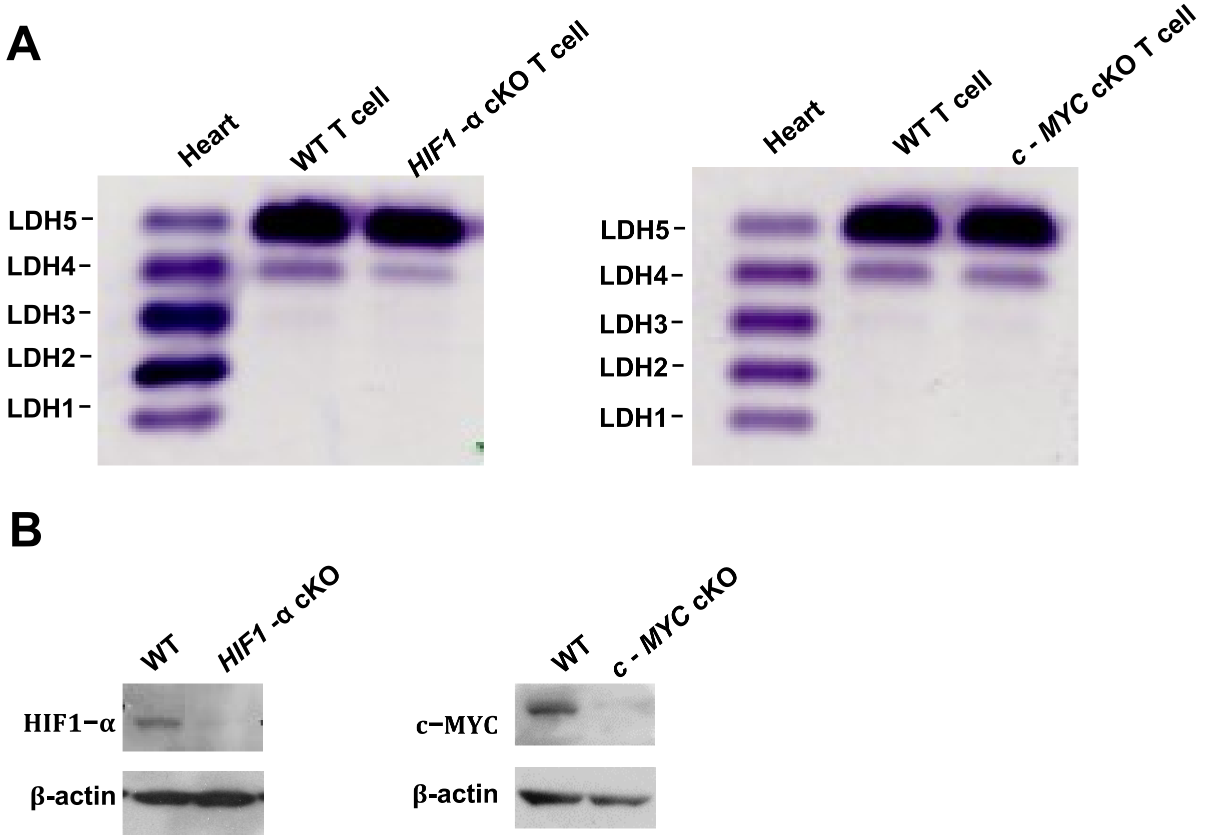


**Supplemental Figure 2. Transcription factors HIF1 and Myc are dispensable for regulating the LDH isoenzyme spectrum in T cells.**

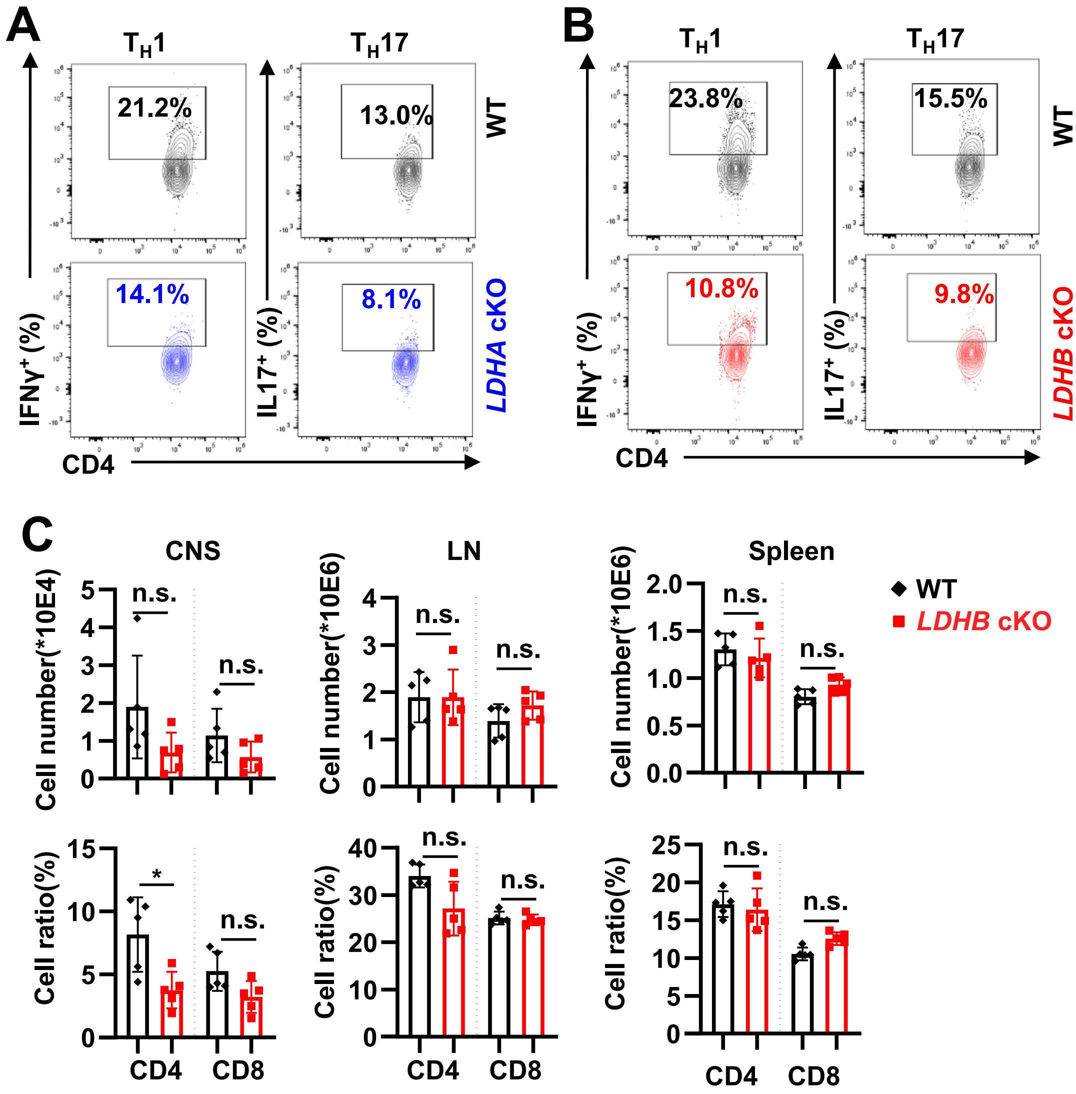


**Supplemental Figure 3. Alteration of LDH isoenzyme spectrum impacts T cell differentiation in vivo.**

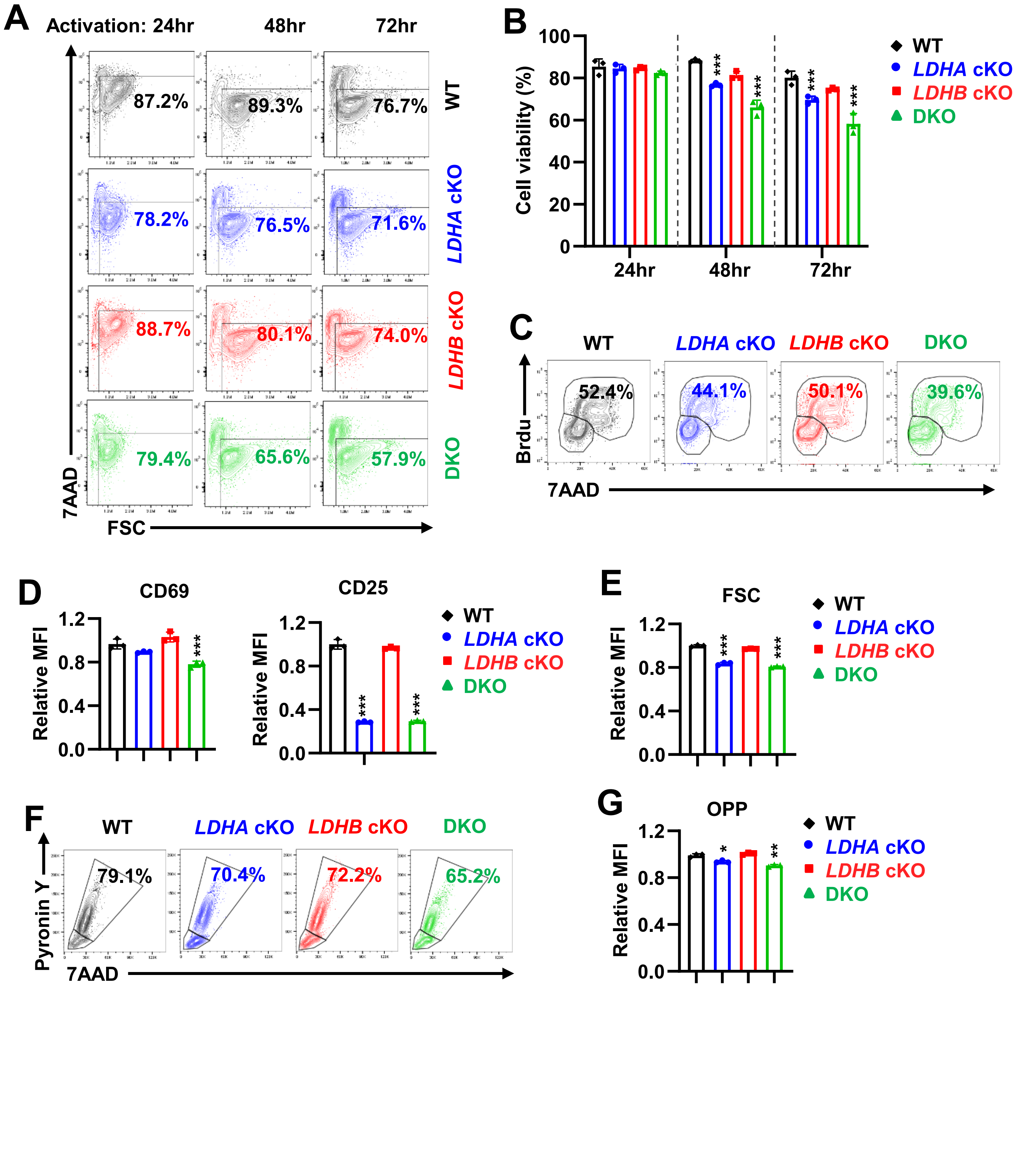


**Supplemental Figure 4. Alteration of the LDH isoenzyme spectrum differentially impacts T cell activation in vitro.**

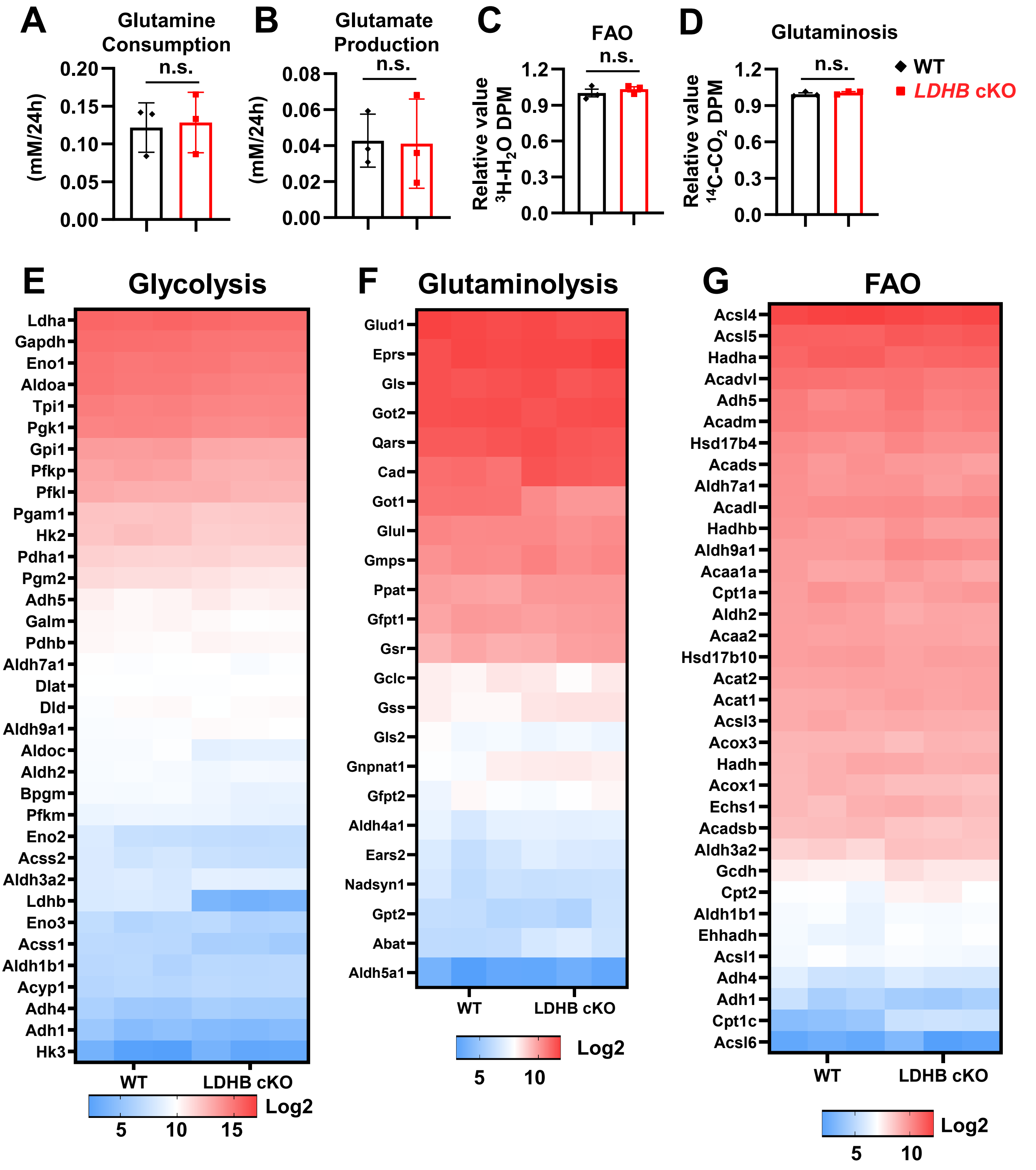


**Supplemental Figure 5. Alteration of the isoenzyme spectrum by LDHB deletion does not affect glutamine and fatty acid catabolism.**

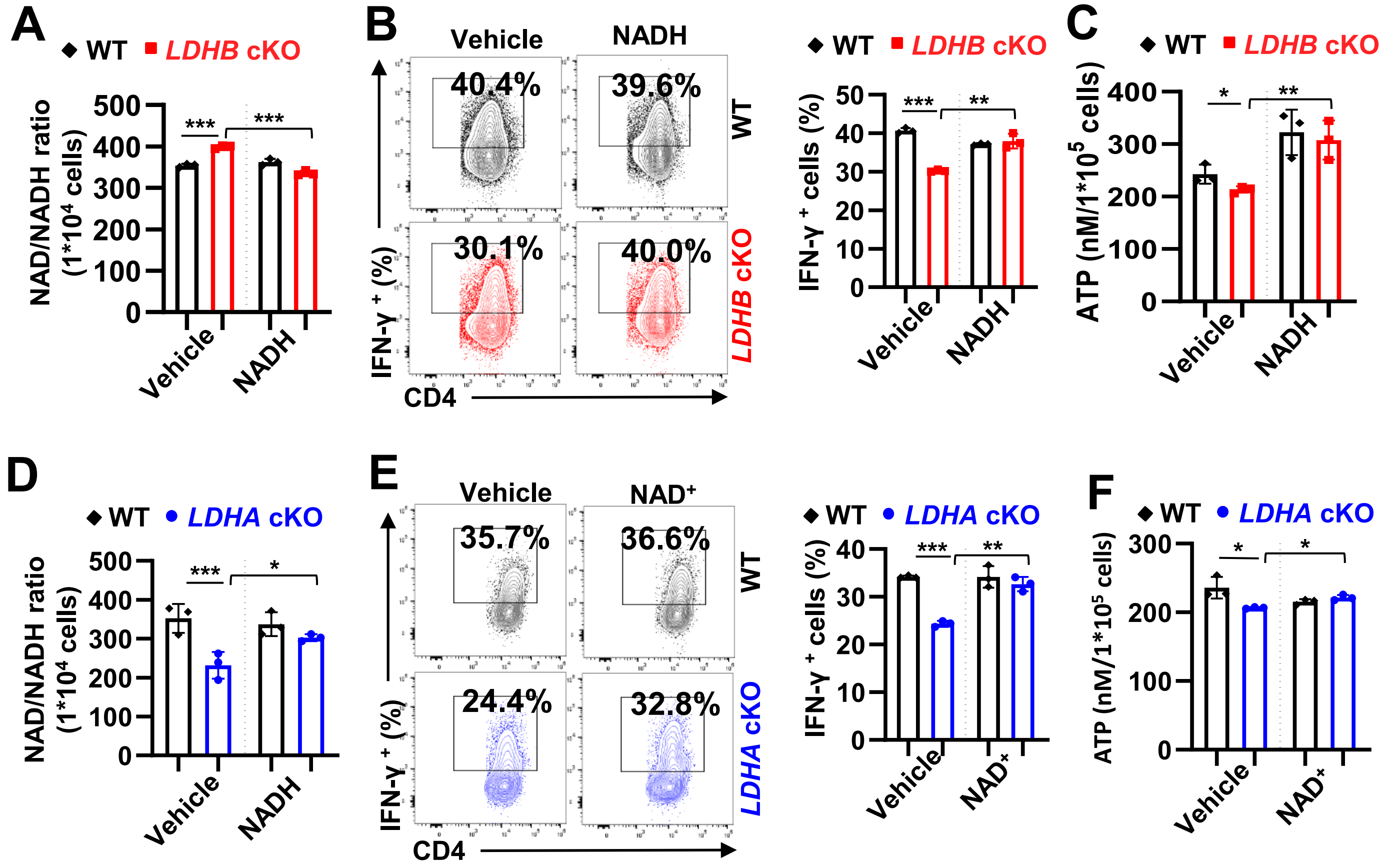


**Supplemental Figure 6. The LDH isoenzyme spectrum confers an optimal glycolytic flux to maintain the NAD⁺/NADH ratio during T cell differentiation.**
