## Supplementary material for "The Goldilocks Principle conferred by LDH isoenzymes controls murine T cell glycolysis and differentiation": table

**Table 1. Cell culture related antibodies, cytokines, and chemicals**

| Name | Cat# | Vendor |
|---|---|---|
| InVivoMAb anti-mouse CD3 | BE0001-1 | BioXcell |
| InVivoMAb anti-mouse CD28 | BE0015-1 | BioXcell |
| InVivoMAb anti-mouse IFNγ | BE0055 | BioXcell |
| InVivoMAb anti-mouse IL-4 | BE0045 | BioXcell |
| InVivoMAb anti-mouse IL-2 | BE0043 | BioXcell |
| Recombinant Murine IL-12 p70 | 210-12 | Peprotech |
| Recombinant Mouse IL-2 | 212-12 | Peprotech |
| Recombinant Murine IL-7 | 217-17 | Peprotech |
| Recombinant Murine IL-6 | 216-16 | Peprotech |
| Recombinant Human TGF-β1 | 100-21C | Peprotech |
| NADH disodium salt | 10107735001 | Sigma |
| NAD+ | S2518 | Selleckchem |

**Table 2. Cell staining antibodies and dyes**

| Name | Cat# | Vendor |
|---|---|---|
| Pacific Blue™ anti-mouse CD4 Antibody | 100428 | BioLegend |
| APC anti-mouse CD4 Antibody | 100516 | BioLegend |
| PE Rat Anti-Mouse CD4 Antibody | 553730 | BD Pharmingen™ |
| APC/Cyanine7 anti-mouse CD8a Antibody | 100714 | BioLegend |
| APC anti-mouse TCR β chain Antibody | 109211 | BioLegend |
| PE/Cyanine7 anti-mouse CD90.1 (Thy-1.1) Antibody | 202518 | BioLegend |
| APC/Cyanine7 anti-mouse CD90.2 Antibody | 105328 | BioLegend |
| APC anti-mouse CD90.2 (Thy-1.2) Antibody | 140311 | BioLegend |
| PE/Cyanine7 anti-mouse CD45.1 Antibody | 110730 | BioLegend |
| APC/Cyanine7 anti-mouse CD45.2 Antibody | 109824 | BioLegend |
| FITC anti-mouse CD45.2 Antibody | 109806 | BioLegend |
| PE-Cy™7 Hamster Anti-Mouse CD69 | 552879 | BD Pharmingen™ |
| CD25 Monoclonal Antibody (PC61.5), PE | 12-0251 | eBioscience™ |
| APC anti-mouse IFN-γ Antibody | 505810 | BioLegend |
| APC anti-mouse IL-17A Antibody | 506916 | BioLegend |
| Pyronin Y | 92-32-0 | Sigma-Aldrich |
| APC anti-BrdU Antibody | 364114 | BioLegend |
| 7-AAD Viability Staining Solution | 420404 | BioLegend |

**Table 3. Immunoblot antibodies**

| Name | Cat# | Vendor |
|---|---|---|
| LDHB Antibody | NBP1-55415 | Novus |
| HIF-1α Antibody | 10006421 | Cayman |
| c-Myc Antibody | 9402S | Cell Signaling |
| Anti-actin Antibody | SC-47778 | Santa Cruz |

**Table 4. RT-qPCR primers**

| Gene | primer sequences forward | primer sequences reverse |
|---|---|---|
| Ldha | CATTGTCAAGTACAGTCCACACT | TTCCAATTACTCGGTTTTTGGGA |
| Ldhb | GGGAGCTTGTTCCTCCAGAC | TGGGTTGGAAACCACGATGAT |
| Tubulin | TTCTGGTGCTTGTCTCACTGA | CAGTATGTTCGGCTTCCCATTC |
